# Nuclear lamina-associated domain biogenesis is regulated by nuclear pore density during embryogenesis and mediates UV protection

**DOI:** 10.1101/2025.10.01.679595

**Authors:** Fei Xu, Adrián Fragoso-Luna, Itai Sharon, Angelo L. Angonezi, Ohad Medalia, Peter Askjaer, Susan E. Mango

## Abstract

Lamina-associated domains (LADs) are critical for genome organization and function, but their formation during development is not well understood. Here, we use DamID and image analysis to reveal the dynamics of LAD biogenesis during *C. elegans* embryogenesis. At early stages, DNA at the lamina is transcriptionally active and lacks lamina-associated heterochromatin. This state depends on abundant nuclear pores, which prevent heterochromatin accumulation at the nuclear periphery. As development proceeds, pore numbers decline, enabling heterochromatin to access the lamina. Reducing nuclear-pore components induces precocious accumulation of heterochromatin to the lamina. Functionally, we find that heterochromatic LADs confer protection against ultraviolet (UV) radiation. Older embryos are resistant to UV light and concentrate DNA damage at the nuclear periphery, whereas early embryos are UV sensitive and accumulate damage throughout nuclei when unshielded by their mothers *in utero*. These findings identify embryonic dissipation of nuclear pores as a key step in heterochromatic LAD assembly, allowing older embryos to withstand exposure to UV irradiation.

## MAIN

In animal nuclei, genomic DNA is organized into distinct spatial domains that contribute to their regulation and function. One critically important domain for nuclear organization is the lamina-associated domain (LAD), which comprises genomic regions localized to the nuclear periphery^1^. LADs are classically associated with transcriptional repression and enriched in densely packed heterochromatin that is marked by repressive histone modifications such as histone H3 methylated on lysine 9 (H3K9me3)^1–3^. Disruption or loss of LADs has been linked to aging and cancer, revealing their critical role for cellular vigor^1^. Although positioning of heterochromatin at the nuclear lamina has been observed for decades, little is known about the timing and regulation that establish LADs during development. Recent studies with mice have shown that LADs are established *de novo*, with increasing genomic coverage during embryogenesis and with input from histone modifications^4–7^.

To identify lamina-associated loci in early *C. elegans* embryos, we fused DNA adenine methyltransferase (Dam) to the nuclear envelope protein EMR-1/emerin and expressed the chimera in a subset of embryonic cells prior to the onset of gastrulation (Fig. 1a)^8–10^. Specifically, we chose the EMS mesendodermal lineage from the 4-28-cell stage, to provide a homogeneous cohort of cells for analysis. Single-molecule RNA FISH (smFISH) confirmed expression of the fusion between the 4-cell and 28-cell stages (**Extended Data** Fig. 1a). The fusion strain was healthy and developed normally; DamID replicates were highly reproducible (**Extended Data** Fig. 1b).

**Fig. 1:**
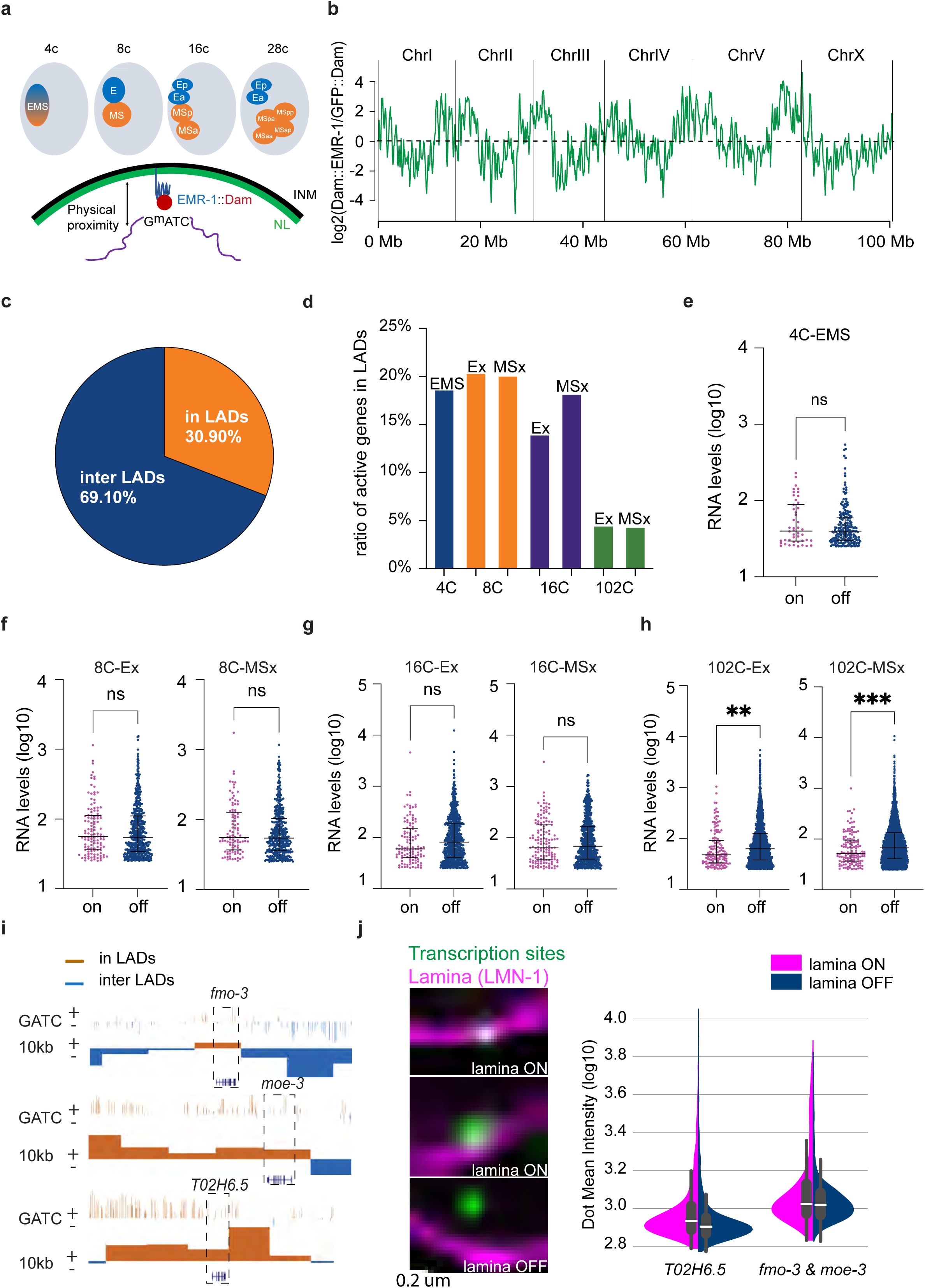
The nuclear lamina is transcriptionally permissive in early embryos. **a,** Schematic of early DamID profiling in the EMS lineage. **b,** DamID signal across the genome shows enrichment in chromosome ‘arms’. **c,** the proportion of genes enriched on or off the lamina. **d,** proportion of actively transcribed genes on the lamina at each indicated cell stage. **e-h,** scRNA-seq expression levels of active genes on vs off the lamina at the indicated cell stage. Each dot represents a gene. Lines indicate median with interquartile range. p (**) < 0.01, p (***) < 0.001 by Mann-Whitney U test. **i,** selected lamina-associated genes showing enrichment (brown) or depletion (blue) of DamID signal (GATC). UCSC genome browser view of reads mapped to GATC sites and enriched 10 kb bins are shown. **j,** images of smFISH signal (green) for introns located on vs. off the lamina (pink) for *T02H6.5*. Violin plot smFISH intensity of indicated genes on (pink) vs off (blue) the lamina. The line represents the median value. Box plots show the interquartile range with whiskers representing minimum and maximum values within IQR. N=2 biological replicates; >100 cells for each gene, per replica. RNA data from^1,2^.

Lamina-associated domains from early embryos exhibited a typical arm-center-arm organization that resembled the profiles of late LADs in *C. elegans*^11–13^. Approximately 31% of genes were located at the lamina in early-stage embryos, which was lower than the 46% in older, mixed-stage embryos, indicating that LAD coverage increases over time (Fig. 1b and **Extended Data** Fig. 1c)^12,13^. To assess the transcriptional activity of these genes, we examined single-cell RNA datasets for genes expressed zygotically both on and off the lamina^14–16^. To do so, we removed the cohort of maternally-expressed RNAs that are detected prior to ZGA at the 4-cell stage, and focused on RNAs induced after the 4-cell stage. This analysis revealed that up to the 16-cell stage, approximately 20% of lamina-associated genes were actively transcribed (Fig. 1c), with expression levels comparable to genes off the lamina (Fig. 1d–f and **Extended Data** Fig. 1d). By the 102-cell stage, both the fraction of expressed lamina-associated genes (Fig. 1c) and their expression levels (Fig. 1g) decreased, indicating that the lamina was becoming a transcriptionally repressive domain during gastrulation.

Single-molecule RNA FISH (smFISH) using intron probes targeting genes in early LADs confirmed active transcription at the lamina in early cells (Fig 1j). DNA FISH combined with smFISH for RNA showed no significant difference in the distribution between transcribing and silent DNA loci, supporting the notion that position did not correlate with transcriptional status at early stages (**Extended Data** Fig. 1e). At later stages, the transcriptional foci shifted towards the interior (**Extended Data** Fig. 1f) in agreement with prior studies^17^. These findings reveal that early LADs are transcriptionally permissive in early *C. elegans* embryos, and that they initiate silencing during gastrulation, starting by the 100-cell stage.

In differentiated cells, LADs are heterochromatin domains, enriched for methylated histone H3K9. To track the emergence of heterochromatin during development, we stained embryos for H3K9me2 and H3K9me3, which are markers for heterochromatin and lamina association ^11,18–20^. H3K9me2 was virtually undetectable in the youngest embryos, consistent with previous studies ^21,22^. Reproducible H3K9me2 signal appeared around the 25-cell stage and began to concentrate at the lamina by the 100-cell stage (Fig. 2a, b). During this transition, nuclei displayed a mottled pattern, with discrete H3K9me2 puncta at the lamina, but extensive sections of the lamina still unoccupied until late stages (Fig 2a, b). H3K9me3 and the MET-2/SETDB methyltransferase followed a similar spatial pattern, whereas H3K27me3 was distributed throughout the nucleoplasm with only modest lamina enrichment, underscoring the distinct distributions of histone marks associated with constitutive versus facultative heterochromatin (**Extended Data** Fig. 2a and 2b). These data reveal that heterochromatin accumulates at the lamina gradually, forming puncta that do not fully coalesce into a continuous peripheral layer until late embryogenesis. We detected transcriptional activity in regions unoccupied by H3K9me3, as monitored by smFISH for lamina-associated genes or for elongating RNA Polymerase II (Fig 2b, c and **Extended Data** Fig. 2c). Active transcription was rarely detected within H3K9me3-enriched domains, suggesting that lamina-associated heterochromatin accounts for most gene silencing, whereas lamina-associated chromatin that is free of heterochromatin remains transcriptionally permissive even in late embryos. This idea is supported by recent screens in human cells that identified heterochromatin factors, but not lamina factors, as drivers of lamina-associated silencing^23^. In short, heterochromatic LADs nucleate and spread across the lamina during embryogenesis and appear to be the principal mechanism of repression.

**Fig. 2:**
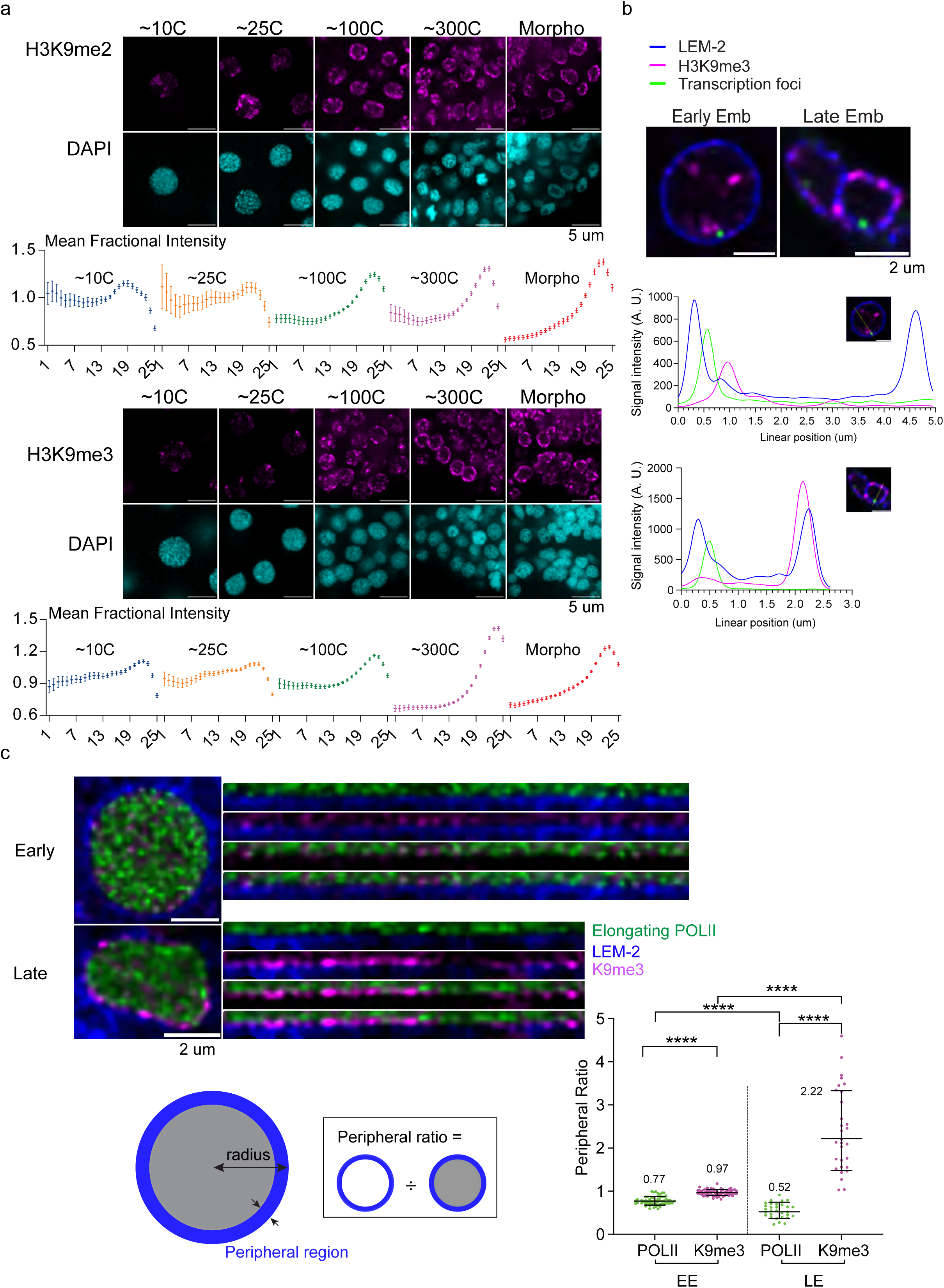
Gradual establishment of the nuclear lamina as a silencing domain. **a,** Images of H3K9me2 and H3K9me3 during development (pink). Plots show the mean fractional intensity across radial regions from the center (1) to the periphery (25) across the x axis. Early to late stages show a gradual polarization of signal towards the periphery. Lines represent mean with SEM. N = 3 biological replicates. **b,** Intron smFISH (green), H3K9me3 (pink) and the lamina (blue) at early and late stages. Line scan shows the spatial relationship for each image. N=3 biological replicates. **c,** Immunostaining of active Pol II (green), lamina (blue) and H3K9me3 (pink) at early and late stages. The right shows a straightened view of the nuclear periphery. Schematic shows the peripheral ratio was calculated from the mean intensity at the periphery (blue) divided by the mean intensity across the entire nucleus (blue + gray). Plots show the peripheral ratio for stains at early and late stages. Each dot shows a nucleus. Lines represent mean with standard deviation. p (****) < 0.0001 by Welch’s t test. N=2 biological replicates.

What delays the association of heterochromatin with the nuclear lamina until gastrulation? Previous studies have suggested an antagonism between nuclear pores and heterochromatin^1,24,25^. Imaging of mouse and human cells displays a strong correlation between heterochromatin-exclusion zones (HEZs) and the presence of nuclear pores. Functional studies with cells under pathogenic or stress conditions have suggested that pores can modulate the location of heterochromatin on or off the lamina^26,27^, raising the question of whether pores could regulate heterochromatic LADs under physiological conditions. To begin to test this idea, we screened embryos for nuclear pore components in developing *C. elegans* embryos. We observed a pronounced enrichment of nuclear pore complex (NPC) proteins during pre-gastrula stages (Fig. 3a, **Extended Data** Fig. 3). Using fluorescently-tagged strains, we analyzed components from the NPC including the nuclear basket (NPP-7/NUP153, NPP-21/TPR), scaffold of the inner ring (NPP-13/NUP93, NPP-19/NUP35), linker (NPP-10N/NUP98), cytoplasmic filaments (NPP-9/NUP358), nuclear ring component (MEL-28/ELYS) and FG repeats of the central channel (mAB414 antibody stains). All showed high expression and a broad distribution across the nuclear envelope at the earliest stages (≤25-cell), but their levels declined as embryos aged, and the signal became punctate. We confirmed this result by transmission electron microscopy (TEM), which revealed abundant pores at early stages and a reduction of more than two-fold in older embryos (Fig. 3b). We note that at least a proportion of pre-gastrula pores are functional since zygotic mRNAs are exported to the cytoplasm and nuclear proteins are imported into the nucleus in pre-gastrula embryos^28–31^. The findings reveal robust accumulation of NPC components at the nuclear periphery in early embryos, followed by gradual reduction of pores during development.

**Fig. 3:**
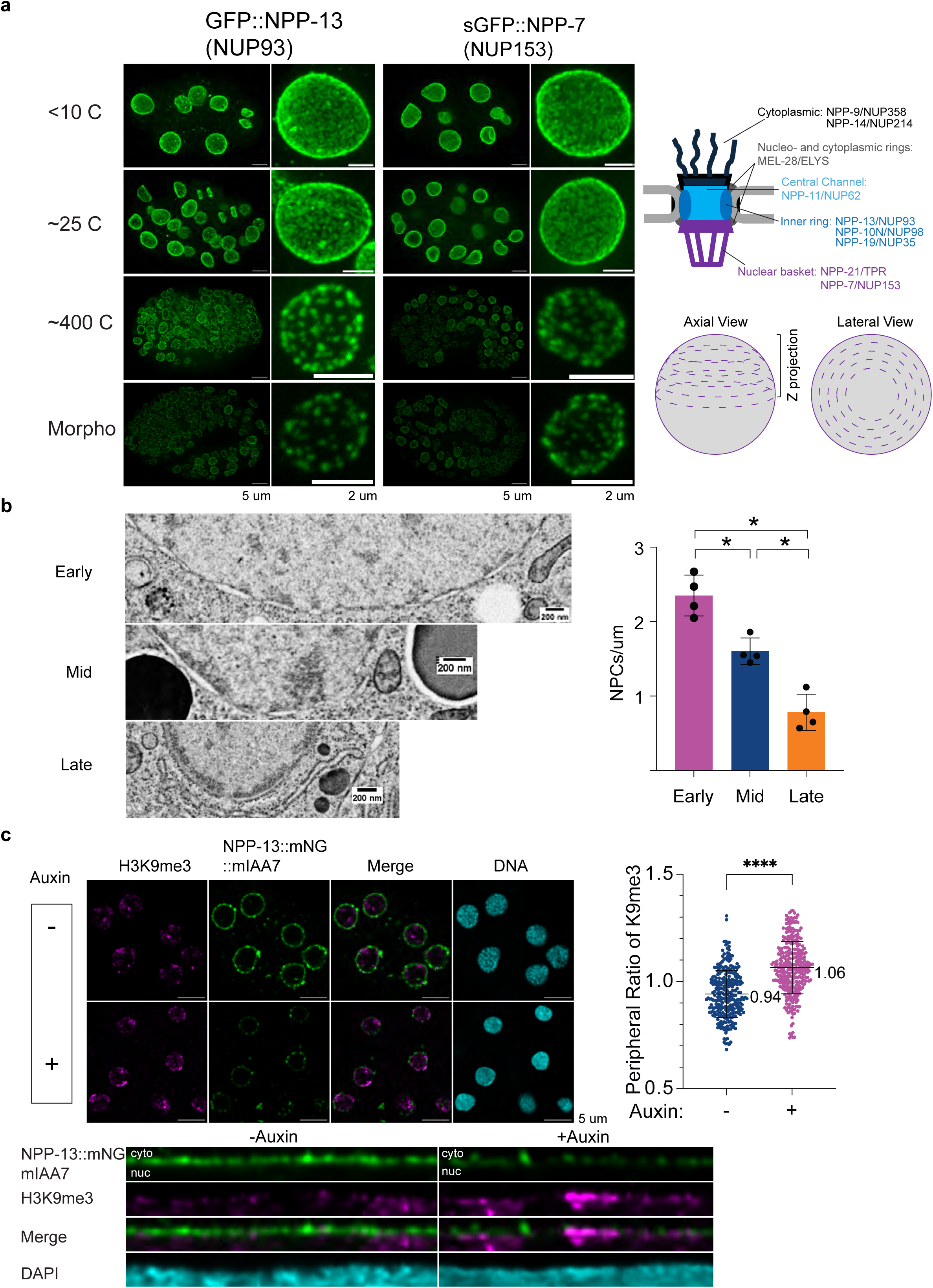
Nuclear pore density regulates the timing of LADs formation. **a,** Immunostaining of the indicated nucleoporins from fewer than 10 cells (<10 C) to morphogenesis. The top-right cartoon illustrates different nuclear pore components. The bottom-right schematic shows the z-projection used to visualize pore density on the surface of half the nucleus. **b,** Electron microscopic images of nuclei during embryogenesis. Bar plots show the number of pores per micron at the indicated stages. Each dot represents one nucleus. Lines represent mean with standard deviation. p (*) < 0.05 by Mann-Whitney U test. **c,** Images show H3K9me3 (pink) before (-) and after (-) 15-minute auxin-induced degradation of NPP-13/NUP93 (green). The bottom images show a straightened view of the nuclear periphery. The dot plot compares the peripheral ratio of H3K9me3 with and without 15 minutes Auxin-induced degradation. Each dot represents a nucleus. Lines indicate mean with SD. p (****) < 0.0001 by Welch’s t test. N=4 biological replicates, total ≥283 cells with or without auxin.

We hypothesized that the abundance of pore proteins in early embryos might interfere with the formation of heterochromatin LADs. Consistent with this idea, regions with pores, observed by TEM, lacked electron-dense, nascent heterochromatin (Fig 3b). Similarly, stains for the nuclear pore components NPP-13::mNeonGreen/NUP93 or NPP-21::GFP/TPR revealed minimal colocalization with H3K9me3 (Fig. 3c and **Extended Data** Fig. 4d). These observations suggested that the high concentration of nuclear pores in early embryos precluded the accumulation of heterochromatin at the lamina. To test this idea, we partially depleted the inner ring protein NPP-13/NUP93 (*npp-13(LF*)) by auxin-induced degradation (AID) system or RNAi^32,33^. The goal was to reduce the number of pores, while leaving some intact to carry out necessary biological functions. 15-minute treatment of AID-tagged NPP-13 in early embryos reduced its abundance to 45% of the control. 30- to 45-minutes of treatment further decreased NPP-13 levels to approximately 25% of the control (**Extended Data** Fig. 4a). Partial depletion of NPP-13 led to a reduction of FG repeat-containing nucleoporins at the nuclear periphery, indicating pores were disrupted (**Extended Data** Fig. 4b). As a complementary approach, we also performed longer-term, partial reductions using RNAi-treated worms with a diluted *npp-13* double-stranded RNA, leading to reduction of NPP-21::GFP/TPR signal at the periphery (**Extended Data** Fig. 4d). This is consistent with the notion that scaffold protein NPP-13/NUP is required for proper assembly of nuclear pores^34,35^.

We performed three controls that showed cells with reduced pore components were still functional. First, treated cells were permissive for RNA Pol II transcription, as measured by immunofluorescence for phosphorylated, elongating RNA Pol II (**Extended Data** Fig. 4c). Second, we observed the proper distribution of zygotic transcriptional factor ELT-2 in E/gut cells, indicating that *elt-2* RNA was exported after transcription, and nuclear ELT-2 protein was imported after translation, as expected (**Extended Data** Fig. 4e). The accumulation of ELT-2 protein was decreased somewhat, consistent with a decreased number of pores (**Extended Data** Fig. 4e). Third, we observed nuclear MET-2/SETDB1 and HPL-2/HP1, indicating that components of the heterochromatin machinery were properly localized in nuclei, despite fewer pores (**Extended Data** Fig. 4f and 4g). These controls indicate that cells retained active transcription and transport despite reduced accumulation of nuclear pore proteins.

Reduction of *npp-13*/NUP93 by AID-mediated degradation increased perinuclear H3K9me3 in pre-gastrula embryos. We observed crescents and blobs of H3K9me3 at the lamina already by the 10-cell stage, a stage when heterochromatin never accumulates at the lamina in the wild type. We did not see full coverage of the nuclear periphery after 15 minutes auxin, as is observed in mid- or late-staged wildtype embryos, but longer treatment (e.g. 45 minutes) produced greater coverage compared to 15 minutes depletion (Fig. 3c, **Extended Data** 4a, d). As in untreated embryos, H3K9me3 was visible in lamina regions lacking NPP-13::mNeonGreen or NPP-21::GFP, but absent from regions that retained strong pore signals (Fig. 3c and **Extended Data** Fig. 4d). This result suggests that, normally, pore components delay heterochromatin-lamina interactions until after gastrulation, when pore density declines. The converse was not true: removal of virtually all H3K9me in *met-2; set-25* double mutants^11^ did not disrupt the distribution or levels of pore components in developing embryos at early or late stages (**Extended Data** Fig. 4h, i). Together, these data suggest that the lamina of early embryos is transcriptionally active because abundant nuclear pores impede the formation of lamina-associated heterochromatin.

Next, we addressed the purpose of lamina-associated heterochromatin that is present in older but not younger embryos. One proposal is that a shell of peripheral heterochromatin shields nuclei from exogenous damage such as ultraviolet light^36–38^. To test this idea for *C. elegans* embryos, we first determined that UVB light at 295 nm was the most effective wavelength for inducing damage in the embryo, as evidenced by increased embryonic lethality (**Extended Data** Fig. 5a). Because DNA is highly mobile within nuclei, especially when undergoing repair^39^, we gently fixed nuclei prior to UVB irradiation. This approach allowed us to monitor where damage occurred. Using these conditions, UV damage was dispersed throughout nuclei in early embryos, but was enriched at the nuclear periphery in late embryos (Fig. 4a). In addition, there was considerably more damage in early embryos compared to late. The amount of UV damage did not correlate with local DNA abundance, supporting the idea that LADs serve as protective shell rather than merely reflecting an increased concentration of DNA at the lamina (Fig. 4a).

**Fig. 4:**
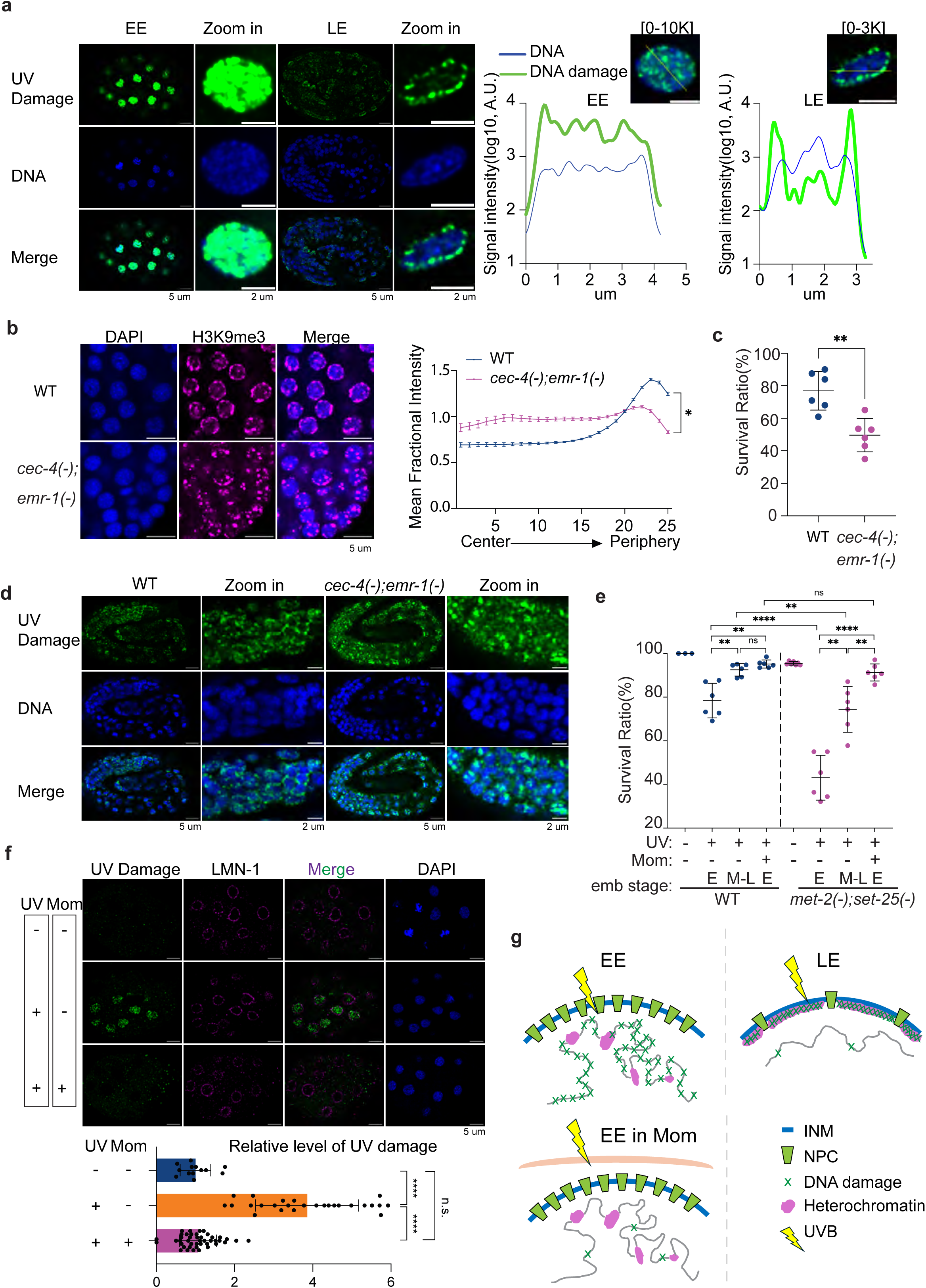
LADs shield nuclei against UV damage. **a,** Images of UV damage (green) in early (EE) and late (LE) embryos. Line scan shows the spatial relationship between UV damage (green) and DNA (blue). Different maximum intensity scales (0-10K, 0-3K) were used to highlight distribution details in the early embryo. N = 3 biological replicates. **b,** Immunostaining of H3K9me3 (pink) in wild-type (WT) and mutant embryos. Plot shows the mean fractional intensity across radial regions. p (*) < 0.05 by Wilcoxon test. N = 3 biological replicates **c,** Embryo survival after UV illumination. Each dot represents one dish, >50 embryos per dish. Lines represent mean ± standard deviation. p (**) < 0.01 by Welch’s test. N = 3 biological replicates **d,** Images of UV damage in wild type (WT) and mutant embryos. N = 3 biological replicates. **e,** Proportion of surviving embryos after UV illumination for early (E) or mid-late (M-L) embryos. Embryos were located in (+) or outside (-) of mothers. Each dot represents a culture dish, >100 embryos per dish. Lines represent mean with SD. p (**) < 0.01, p (****) < 0.0001 by Welch’s test. N = 3 biological replicates. **f,** images of UV damage (green) in early embryos in (+) or outside (-) of mothers. Plot shows UV damage normalized to mean of non-UV control group. Each dot represents an embryo. Lines represent mean with SD. p (****) < 0.0001, p (ns) > 0.05 by Welch’s test. N = 2 biological replicates. **g,** working model: LADs in late embryos or mothers protect DNA within the nuclear interior from UV damage, and this is timed by the density of nuclear pores.

To probe the role of lamina-associated heterochromatin, we searched for conditions that removed heterochromatin from the lamina but kept H3K9me intact. The perinuclear-localized chromodomain-protein CEC-4 targets H3K9me in repeat-rich regions to the lamina, while the LEM-domain-containing protein EMR-1/emerin anchors unique genes to the lamina^40,41^. A double mutant of *cec-4; emr-1* displayed disrupted LADs, with H3K9me3 signal displaced from the periphery (Fig. 4b). Therefore, the double mutant provided a condition with intact, but mis-localized heterochromatin. Upon UVB irradiation, *cec-4; emr-1* mutants had reduced survival and frequent CPD staining in the nucleoplasm (Fig. 4c, d). Late-stage, wildtype embryos survived and displayed an enrichment of mutated cyclobutene pyrimidine dimers (CPDs) at the nuclear periphery (Fig. 4bc, d). We conclude that LADs help animals survive UV irradiation by focusing damage at peripheral heterochromatin. Consistent with this idea, disruption of heterochromatin in a *met-2; set-25* double mutant was very sensitive to UVB light (Fig. 4e).

Finally, we examined young embryos, which lack heterochromatic LADs and which are contained within mothers until gastrulation, when LADs begin to assemble. We hypothesized that mothers might shield early-stage embryos from UV damage. Consistent with this idea, when removed from their mothers, early embryos were highly sensitive to UVB treatment. We detected CPD staining throughout nuclei and poor survival (Fig 4e, f). Within mothers, however, early embryos survived exposure to UVB and exhibited considerably less damage than their freed counterparts (Fig 4f). When young *met-2; set-25* mutants were subjected to UV exposure within mothers, their survival resembled that of the wild type, suggesting maternal protection of UV exposure compensated for the loss of heterochromatin (Fig. 4e). These data also imply that the reduced survival of *met-2; set-25* mutant embryos did not reflect aberrant transcription or disrupted heterochromatin *per se,* as these would not be suppressed by the presence of the mother. We suggest that mothers shield their progeny from radiation at early stages, giving embryos time to build a protective shell of heterochromatin as they mature.

In summary, these results show that embryos are born with a transcriptionally-permissive nuclear lamina, which gradually transitions into a silencing domain as heterochromatin accumulates to form LADs. We suggest this transition is temporally regulated by nuclear pore complexes, which are abundant in early embryos and exclude heterochromatin from the periphery. The rapid degradation of an inner ring pore component, together with the dynamics of nuclear pores in early embryos^42^, enabled us to test this hypothesis. Our data suggest that as development progresses and pore numbers decline, heterochromatin gains access to the nuclear lamina and spreads across the nuclear periphery. This timing is closely coordinated with egg laying. Early embryos that lack LADs are protected *in utero* by mothers, whereas after egg laying, embryos rely on their nascent LADs. Pores are not the only temporal regulators, as recent studies have suggested additional factors^6,7^, however they are sufficient to delay heterochromatin accumulation at the lamina in pre-gastrula embryos. Nuclear pores associate with highly active genes such as Pol III-transcribed genes, as well as architectural proteins, which may explain why they exclude heterochromatin^35,43–45^. Given the variability in nuclear pore density across cell types^46,47^, we speculate that this mechanism may represent a general strategy for modulating heterochromatic LAD density across diverse cellular contexts.

Although heterochromatin was excluded from the nuclear lamina in early embryos, active chromatin remained capable of associating there. DamID profiling showed that genomic regions destined to become heterochromatic were already in contact with the lamina, and that the overall pattern of lamina-associated domains changed quantitatively rather than qualitatively over time. We did not observe large blocks of heterochromatin associated with small stretches of pore-free lamina but rather heterochromatin concentrated within the nuclear interior. The variability of heterochromatin observed early may explain a conundrum in the field, namely why some genes appear to escape lamina-associated silencing^48^. These cases may reflect discontinuities in heterochromatin deposition that allow lamina-associated genes to be transcriptionally active in the proportion of cells where they embedded in euchromatin. Our single-cell analysis revealed this interspersed pattern of LADs and active loci.

Our findings on UV-induced damage may have broader relevance. UV radiation from sunlight is a well-established source of DNA damage, particularly for skin cancer. Genomic analyses of melanoma cells have revealed an enrichment of C→T transition mutations within LAD-associated, cancer-related genes, indicating a UV damage signature that is evident both in DNA sequence and in spatial genome organization^36^. Our findings suggest that variation in LAD composition and nuclear pore density may underlie differences in cellular susceptibility to UV-induced damage across developmental stages and cell types. Conversely, previous studies discovered that photoreceptor cells in several nocturnal species lack LADs altogether^49^. Nocturnal animals are active when there is little UV, possibly explaining why these animals were able to reorganize their heterochromatin domains without incurring the harmful effects that normally accompany the loss of LADs.

## Acknowledgements

We thank the Biozentrum Imaging Core Facility (IMCF) for technical. Specifically, Alexia Loynton-Ferrand for help in setting up the SoRa super-resolution module. Kai Schleicher assisted with Huygens deconvolution and the design of the stage insert for the Ti2 microscope. Laurent Guerard and Sébastien Herbert for help in imaging analysis. We thank Lionel PINTARD’s lab for sharing the transgenic strains of nucleoporin proteins used in this study. Morris Maduro for advice on the med-1 promoter as well as Cristina Ayuso, Beatriz Ren and Ildefonso Cases for technical assistance. Cristina Tocchini for technical assistance in constructing transgenic worms. Michael Steinacher from the Department of Physics provided assistance with UV measurements. Oded Mayseless for sharing the UVA illumination LED. The Central Mechanical Workshop for making the stage insert for the Ti2 system. Christian Oxé from the Central Electronic Workshop developed the UVB and UVC illumination systems. WormBase for resources. A. Schier for reading a draft of the manuscript. Some strains were provided by the CGC, which is funded by NIH Office of Research Infrastructure Programs (P40 OD010440). Some calculations were performed at sciCORE (http://scicore.unibas.ch/) scientific computing center at University of Basel. Funding was SNSF 320030-227954 to SEM; SNSF grant 320030L-231406 to OM, the Spanish Ministerio de Ciencia, Innovación y Universidades (MCIU), Agencia Estatal de Investigación, the European Union and the European Regional Development Fund (CEX2020-001088-M, and PID2022-137162NB-I00; doi:10.13039/501100011033).

## Material and Methods

Strain maintenance: All strains were maintained at 20°C.

DamID:

Molecular cloning^50^

Plasmids pBN560 med-1p::dam::emr-1 and pBN561 med-1p::gfp::dam for DamID were constructed by Gibson assembly inserting a gBlock containing a 436 bp med-1 promoter (position -436 to -1 relative to the med-1 AUG start codon) and flanking sequences into pBN209 hsp-16.41p>mCh::his-58>dam::emr-1 or pBN181 hsp-16.41p>mCh::his-58>gfp::dam, respectively, cut with AvrII and XhoI.

### Mos1-mediated single-copy integration

Single-copy insertion of the DamID constructs into the ttTi5605 locus on chrII was done by microinjection into the gonads of EG4322 young adult hermaphrodites. Plasmids pBN560 and pBN561 carrying unc-119(+) as selection marker were injected at 50 ng/µl together with pCFJ601 eft-3p::Mos1^50^ (50 ng/µl), pBN1^51^ (10 ng/µl), pCFJ90^50^ (2.5 ng/µl) and pCFJ104^50^ (5 ng/µl). Integration of the constructs was confirmed by PCR.

### Nematode culture for DamID

Worms were grown asynchronically at 20°C on NGM plates seeded with E. coli GM119 dam-for at least two generations before egg-prepping with hypochlorite. Hatched L1s were counted the day after and 1000 L1s were aliquoted onto 10 cm NGM plates seeded with E. coli GM119. Worms were grown at 20°C for 72 h before collecting 4000 young adults per strain and washed with 15 ml of M9 with 0.01% Tween-20. Worms were transferred to 1.5 ml tubes with M9 with 0.01% Tween-20 using Pasteur pipettes followed by 2 additional washes. Eggs were released by hypochlorite treatment and washed 5 times with M9 with 0.01% Tween-20. Supernatants were aspirated and aliquots of 30 µl were snap frozen in liquid nitrogen and maintained at -80°C until DNA extraction. Three independent biological replicas were prepared.

### DamID amplification, library preparation and sequencing

To purify genomic DNA, samples were lysed in 5 freeze/thaw cycles (1min in liquid nitrogen; 3min in 37C shaker) and processed with DNeasy Blood and Tissue Kit (QIAGEN #69504). DamID was performed on 400 ng genomic DNA with a pool of 3 AdR primers (B1436 AdR4N, B1437 AdR5N, B1438 AdR6N) and 24 PCR cycles as described^52^. Successful amplification of Dam-methylated genomic DNA yielded a smear of ∼400-∼1500 bp fragments when analyzed on Agilent Bioanalyzer.

40 ng of PCR amplified fragments were used for library preparation as described^52^ using E7370 NEBNext® Ultra™ DNA Library Prep Kit for Illumina. Library amplicons’ size distribution was checked to be maintained as for PCR fragments indicated above followed by pooling, repurification and sequencing on an Illumina NextSeq500 platform at EMBL GeneCore.

### Bioinformatic analysis

FASTQ files from the NextSeq500 platform were processed in R Studio using the pipeline RDamIDSeq (https://github.com/damidseq/RDamIDSeq)^53^ for mapping of reads to GATC fragments and to bins of fixed size (e.g. 2 kb, 10 kb or 100 kb). The pipelines include a quality check where only reads that contain the DamID adapter (“CGCGGCCGAG”) and map to GATC sites in the C. elegans genome are retained for further analysis. The number of reads fulfilling these criteria ranged from 15.8 to 22.8 million per sample (see Supplementary Table S1). UCSC genome version ce11 (BSgenome.Celegans.UCSC.ce11) was used for alignment.

The DESeq2 package^54^ was used to identify bins with significantly more reads with Dam::EMR-1 compared to GFP::Dam across the 3 replicas (“bound”). Conversely, bins with significantly more reads with GFP::Dam compared to Dam::EMR-1 were scored as “depleted”. A false discovery rate (FDR) of 0.05 was used as threshold. Next, for each sample a pseudocount of 1 was added to each bin and the relative number of reads per bin was calculated followed by averaging across the replicas. The ratio of Dam::EMR-1 to GFP::Dam was calculated for each bin and log2-transformed. Finally, normalization was done by subtracting the genome-wide average log2 ratio from the log2 ratio of each bin^55^.

Genome browser views were generated with the University of California Santa Cruz Genomics Institute Genome Browser (http://genome.ucsc.edu)^56^.

Note, we first attempted to obtain cell type specific DamID using recombination^57^ or Targeted DamID^58^. However, no recombination activity was observed from multiple med-1p::FLP lines nor were Targeted DamID mRNA robustly detected by FISH. Based on this result and available single-cell RNA expression data, we inferred that the activity of the med-1 promoter is transient and low and therefore compatible with direct expression of dam::emr-1 and gfp::dam mRNAs.

RNAi:

Feeding RNAi was performed as previously described^33,59^. The npp-13 RNAi vector was cloned from C. elegans cDNA and assembled into L4440 plasmid and transformed into HT115 bacteria. RNAi bacteria were grown in LB with 100 ug/ml ampicillin and tetracycline 12.5 ug/ml overnight at 37°C followed by induction with 2 mM IPTG 1 hour. 400 ul of bacteria were then seeded onto 6-cm RNAi plate containing 2 mM IPTG. Seeded plates were then dried and kept at room temperature for 24 hours to induce dsRNA production at room temperature. L4-stage worms were transferred to the plates and incubated at 20°C for 24 hours before dissection. RNAi efficiency was assessed by observing either approximately 100% embryonic lethality at early-to-mid stages following full npp-13 knockdown or the reduction of NPP-21::GFP signal at the nuclear periphery after diluted RNAi treatment.

### smFISH probes

smFISH probes were designed using Oligostan.r^60^ with inputting intron- or exon-sequences based on the detection. Probe sequences were further BLASTed to avoid off-targeting binding especially in protein-coding regions. Probes were synthesized by Integrated DNA Technologies.

### DNA FISH probes

86 probes with length 98 nt were designed to cover the 5kb region [ChrV: 20749945 to 20754904] around the pha-4 promoter. Probe design was based on previously described methods^61^. DNA sequences were extracted from the C. elegans genome assembly (ce11)^62^. Each primary probe sequence contains Genome homologous sequence (42 nt) flanking with readout sequence (28 nt). The OligoArray 2.1^63^ parameters used were: targeting region length 42 nt, melting temperature 78-100°C, no internal secondary structure with melting temperature greater than 76°C, no cross-hybridization with melting temperature greater than 76°C. GC content 30%–90%, and sequences do not contain consecutive repeats of seven or more identical nucleotides. OligoArray2.1 was run on sciCORE for high-performance computing at University of Basel. The 42 nt homologous sequences were further blasted by NCBI BLAST+ 2.9.0 for specificity. Oligos were made by Integrated DNA Technologies.

### High pressure freezing and array tomography

Embryos of C. elegans were harvested using the bleach method^64^. Embryos were spun down and pipetted to the 100 µm cavity of a 3 mm aluminum specimen carrier (Engineering Office M. Wohlwend GmbH, Sennwald, Switzerland). Excess buffer was blotted using filter paper. Subsequently, a flat 3 mm aluminum specimen carrier was dipped in 1-hexadecene (92%, Merck, Buchs, Switzerland) and added on top. The specimen carrier sandwich was transferred into an EM HPM100 high-pressure freezer and frozen instantly (Leica Microsystems, Heerbrugg, Switzerland). Freeze-substitution was carried out using an AFS2 freeze-substitution unit (Leica Microsystems) with integrated, custom built agitation system in water-free acetone containing 1% OsO4 for 10 hours at -90°C, 7 hours at -60°C, 5 hours at -30°C, 1 hour at 0°C, with transition gradients of 30°C/hour, followed by 1 hour incubation at RT. Samples were rinsed three times with acetone water-free, block-stained with 1% uranyl acetate (Electron Microscopy Sciences, Hatfield, PA, USA) in 20% acetone (in methanol) for 1 hr at 4°C, rinsed with water-free acetone and embedded in Epon/Araldite (Merck, Darmstadt, Germany): 66% in acetone water-free overnight, 100% for 1 hour at RT and polymerized at 60°C for 28 hours.

Next, serial ultrathin sections (70nm) were collected on 10mm x 20mm silicon wafers using an ultramicrotome (Artos 3D, Leica Microsystems) (White and Burden, 2023) and contrasted with Reynolds lead citrate for 7 min. The sections were imaged in an Apreo 2 VS scanning electron microscope using the MAPS software package for automatic serial section recognition and image acquisition (Thermo Fisher Scientific, Eindhoven, The Netherlands). The array tomography workflow includes serial sections recognition, image region definition, autofunctions, and image acquisition^65^. Region of interest was imaged on every section using the OptiPlan mode of the system and the T1 detector with following parameters: Pixel size of 4 nm, pixel dwell time of 7 us, electron high tension of 1.8 keV, beam current of 0.1 nA. Serial section Tiff images were aligned using the plugin TrakEM2 (Cardona et al., 2012) in Fiji^66^. Finally, the nuclear pore complexes in the embryos were identified manually and statistically analyzed.

### Immunostaining

Embryo stains were performed as previously described^67^ Embryos were dissected from gravid adults and placed on poly-L-lysine coated slides, then were fixed using fixation buffer (1xPBS/ 0.05%Triton X-100/ 2%PFA) with gentle squishing (squishing more for late-stage embryos until several embryos on the edge burst) at room temperature for 5 minutes. Then the embryo shell was removed by freeze-cracking and embryo samples were merged into ice-cold methanol for 5 minutes. Embryos were washed once with 1xPBS for 5 minutes, three times in 1xPBS/0.5%TritonX-100 for 15 minutes each, and once with 1xPBS for minutes, all at room temperature. Embryos were blocked using IF blocking buffer (1% Normal Goat Serum in 50 mM Tris-HCl pH 7.5 and 150 mM NaCl solution) for 1 hour at room temperature. Primary antibodies were diluted at 1:300 (unless being specifically addressed) in IF blocking buffer at indicated concentration and incubated at 4°C for overnight. Wash the embryos with 1xPBS three times for 5 minutes each. Incubate embryos with secondary antibodies diluted at 1:500 in IF blocking buffer for 1 hour at room temperature. Embryos were then washed in 1xPBS for three times 5 min each. Nuclei were stained with Nuclei were stained with DAPI (1 mg/ml) at 1:1000 dilution in 2xSSC. Slides were sealed with mounting medium VECTASHIELD.

Immunofluorescence for Cyclobutane Pyrimidine Dimers (CPD)((Clone TDM-2 Cosmo Bio) was modified as previously described^68^. After freeze cracking, submerging in ice-cold methanol and washing once with 1xPBS, fixed embryos were illuminated by UVB for 2 minutes. Embryos were post-fixed by fixation buffer for 10 minutes, washed by 1xPBS once for 5 minutes, permeabilized by 1xPBS/0.5%TritonX-100 three times for 15 minutes each. Then 37% HCl was diluted to 2M in water and applied to the embryos for 30 min at room temperature. Sample was then wash in 1xPBS once for 5 minutes and following the blocking and incubating steps as previously described^68^.

IF with the H5 antibody was performed as previously described^69,70^ with modifications. Embryos were gently squished in 1xPBS and freeze-cracked on dry ice. Embryos were fixed in methanol in -20°C for 1minute and immediately followed by fixation solution (3.7% w/v formaldehyde/ 1.6 mM MgSO4/ 0.8 mM EGTA/ 80 mM HEPES/ 1xPBS) at room temperature for 30 min. Embryos were washed in 1xPBS once for 5 minutes and three times in PBT buffer (1xPBS/ 0.1% TritonX-100, 0.1% BSA) three times for 10 minutes each. Block the sample using PBT buffer for 30 minutes at room temperature. H5 was diluted in PBT buffer at 1:200, applied to embryos and incubated at 4°C for overnight. Embryos were washed in 1xPBS three times for 5 minutes each. Goat anti-mouse IgM Alexa Fluor 555 was diluted at 1:200 dilution in PBT buffer and incubated with embryos for 2 hours at room temperature. Embryos were washed with 1xPBS and sealed with DAPI stained.

### Single-molecule FISH

smFISH was performed according to^67^. Embryos were dissected and placed on poly-L-lysine coated slides, then fixed using fixation buffer (1xPBS/ 0.05%Triton X-100/ 1%PFA) with gentle squishing at room temperature for 5 minutes. The embryo shell was then removed by freeze-cracking, and embryos were transferred into ice-cold methanol for 5 minutes. Embryos were then washed with 1xPBS, three times in 1xPBS/0.5%TritonX-100 for 15 minutes each. Embryos were then blocked in hybridization buffer (10% dextran sulfate (m/v)/ 2X SSC/ 10% formamide) for one hour at 37°C. Meanwhile, smFISH probes were pre-annealed as described^60^. Pre-annealed probes were diluted to the final concentration of 3.2 nM for each single probe and applied to the samples. Samples were incubated at 37°C for 4 hours or overnight. Samples were washed by wash buffer (2xSSC/10% formamide) at room temperature. All reagents were nuclease free.

### smFISH combined with DNA FISH

smFISH was performed as previously described^67^ on poly-L-lysine-coated 40 mm round coverslip. Probes targeting pha-4 introns and exons were barcoded with FLAPY-Atto565 and FLAPY-Atto647, respectively. Samples were post-fixed for 10 minutes. DNA FISH was performed as previously described^61,71^. Primary pha-4 DNA probes were diluted in hybridization buffer (2xSSC/ 0.1%% Tween-20/ 10% dextran sulfate/ 50% formamide) to the final concentration of 3.2 nM for each probe. After finishing hybridization and washing steps, secondary FLAPY-Atto565 was diluted in 2xSSC/25% ethylene carbonate to a final concentration of 8 nM and incubated with samples for 30 min at room temperature. Samples were DAPI stained after washing steps.

### Immuno-staining with smFISH

IF performed as previously described with a modified blocking buffer (1.5 mg/ml ultrapure BSA/ 1xPBS/ 0.5 U/ul RiboLock RNase Inhibitor). Post-fixation was carried out using fixation buffer for 10 minutes, followed by a single wash in 1xPBS for 5 minutes. Then, smFISH was performed on the samples as previous described^67^.

### CRISPR/Cas9 transgenesis

Endogenous npp-13/NUP93 was tagged with glycine-serine linker::mNeonGreen::mIAA7 degron^72^ using CRISPR/Cas9-mediated genome editing, as previously described^73^. The repair template was synthesized as a gBlocks Gene Fragment (IDT) and assembled, with assembly product verified by Sanger sequencing (microsynth^74^). The gRNA sequence was designed using the “Design custom gRNA” tool (IDT). crRNA, Alt-R S.p. HiFi Cas9 Nuclease V3 and Alt-R CRISPR tracrRNA were obtained from IDT. The overhang dsDNA repair template was amplified using Phusion Hot Start Master Mix (Thermo Scientific) with nested primers and subsequently annealed. Microinjections were performed with RNP mix containing Cas protein 250 ng/ul, tracerRNA 100 ng/ul, crRNA 56ng/ul, along with 100 ng/ul pRF4 (rol-6(su1006)) coinjection marker and 400 ng/ul of overhang dsDNA repair template. Integrated line was confirmed through Sanger sequencing and was outcrossed once.

### Auxin treatment

Auxin treatment was performed as previously described^75^. 50 mM of eggshell-permeable auxin analog 5-Ph-IAA-AM (Tocris Bioscience) dissolved in DMSO was diluted in 50% M9 buffer or MilliQ water to a final concentration of 4 mM. 5-Ph-IAA-AM induced NPP-13 degradation in the embryos expressing the improved AID2 system of AtTIR1(F79G). Efficient degradation of NPP-13 degradation was induced with a 15-minute treatment in early embryos (before 25-cell stage). 4 mM treatment of N2 Bristol does not induce a significant developmental defect. The survival ratio with and without Auxin treatment is 91.6% and 93.3%, respectively (n>45 embryos for each).

### UV illumination and survival assay

Home-made devices were built using UVB (295 nm) LED Chip on Board (Boston Electronics, VS5252C48L3-295) and UVC (254 nm) LED Chip (Inolux IN-C35PPCTGU0), enclosed in a 3D-printing box, respectively. The consistency and stability of UV power were confirmed by Photodiode Power Sensor (THORLABS S120VC). All experiments were performed at room temperature. For figure 4a and 4d, fixed embryos were directly exposed to UVB for 2 minutes (27.6 mJ/m2) on slides, then stained. For figure 4c, late embryos before comma stage were specifically selected from NGM plates based on embryonic morphology observed under the dissecting microscope. To analyze maternal protection in figure 4e and 4f, early embryos were either dissected from gravid adults and then exposed to UVB on NGM plates, or they were exposed to UVB within mom on NGM plate before being released. The UVB exposure duration was 15 seconds (3.45 mJ/m2). The Survival ratio was calculated as the number of surviving larvae divided by the total number of Larvae and dead embryos.

### Imaging

All images were acquired using Olympus IX83 equipped with Yokogawa CSU-W1 SoRa Confocal Scanner unit, unless otherwise specified. An Olympus 60X (Oil UPL APO, 1.5 NA) objective and a Hamamatsu ORCA-Fusion sCMOS camara were used for image acquisition. Most images were taken at 20°C, except for the live imaging of MET-2::mNeonGreen, which was performed at a controlled room temperature of 15°C. The axial interval was set to 0.2 um, covering a total depth of 20 to 22 um.

Imaging of smFISH combined with DNA FISH was adapted and performed as previously described^76^ using a Nikon Ti2 system equipped with 100x (CFI Plan APO Lambda 1.45 NA) and Photometrics Prime 95B Camara). The experiment was performed: first conducting smFISH imaging within the flow cell chamber (FCS2, Bioptechs). Then, the chamber was disassembled, and DNA FISH was subsequently performed on the same sample. The round coverslip was then reoriented within the chamber, reassembled, and imaged.

### Imaging analysis

Imaging analysis pipelines were implemented using MATLAB (version R2023b; Mathworks.com), Python 3.9^77^, FIJI^66^ macro. Images were pre-processed to correct chromatic aberration using 0.1 um TetraSpeck (Thermo Fisher, T7279) beads as reference.

For figure 1i, signal intensity among embryos was compared by deconvolving images to reduce background noise and enhance resolution using Huygens professional 24.10 (Huygens-Software). Deconvolution parameters were manually optimized prior to batch processing to ensure consistent processing across all images. Three-dimensional nuclear masks were segmented using Cellpose 2.0^78^ with a retrained model. Nuclear lamina and smFISH spots were identified using a difference-of-Gaussian (DoG) edge detector. Lamina-associated spots were defined based on a cutoff of 1% voxels overlapping in 3D.

For figure S1e, smFISH images were aligned to DNA FISH images based on DAPI channel using Descriptor-based registration (2d/3d) in FIJI. Nuclear segmentation was performed via Cellpose 2.0. The 3D coordinates of smFISH spots and DNA FISH spots were determined by 3D gaussian fitting. Cell type classification was based on nuclear association with >10 mRNA molecules. *pha-4* transcribing (ON) nuclei were determined with at least two *pha-4* intron spots, whereas pharyngeal nuclei lacking intron spots were classified as non-transcribing (OFF). The distance from each spot to the periphery was calculated by the nearest distance from spot to the surface using built-in function of alpha shape in MATLAB. For figure S1f, nuclei were segmented using Cellpose 2.0. 3D spot positions and mean intensities were determined by TrackMate^79^ in FIJI using Laplacian-of-gaussian (LoG) edge detector. Spot intensities were subtracted by the mean intensity of their associated nuclei. 2D density distribution was plotted using R studio.

### Mean Fraction Intensity

Mean Fraction Intensity was quantified using MeasureObjectIntensityDistribution in CellProfiler 4.2.6^80^. Individual 2D nuclei from the central focal plane were cropped manually in FIJI. The radial distribution was segmented into bins ranging from 1 (center) to 25 (periphery). The MeanFrac intensity within each radial bin was calculated by normalizing the total intensity to the area of the bin.

### Peripheral ratio quantification

A batch analysis pipeline was developed using Macro in FIJI. Individual 2D nuclei from the central focal plane were cropped manually. Nuclear boundaries were determined based on intensity thresholding plugins in FIJI following Gaussian blur. Peripheral regions were defined using the Distance Map plugin with a threshold of 375 nm. Peripharal areas were generated by subtracting nuclear boundaries from eroded regions through the image Calculator in FIJI. Mean intensities within specified areas were quantified using Measure plugin.

### Signal intensity quantification

Batch analyzing pipelines were built using Macro in FIJI. Channels were split and Z-projection were generated using maximum intensity projections. Nuclei boundaries were identified based on intensity thresholding, and segmented areas were applied to the channel of antibody staining. For measuring elongating RNA Pol II, mitotic nuclei were excluded using Analyze Particles plugin in FIJI, based on size criteria. Background intensity was determined using Make Inverse selection plugin, and mean background intensity was calculated accordingly. Signal intensities were then normalized by subtracting mean background intensity. For fig 4e, individual 2D nuclei of E cells from the central focal plane were manually cropped and fluorescence intensities were measured.

### Line scan analysis

Line scan was performed using Plot Profile or Multichannel Plot Profile plugin in FIJI. Analysis of lamina-associated genes and their expression levels

Data analysis pipelines were implemented using MATLAB 2023B. Single-cell RNA seq data for embryonic stages with up to 16 cells were download from Gene Expression Omnibus (GSE77944)^15^. Detailed dataset annotations were referenced from the BioProject SRA (SRP070155). Read counts were averaged across replicates of each individual cell type, such as Ex, MSx. Processed data validation was conducted by comparison with the data visualization tool available at http://tintori.bio.unc.edu. The expressing cutoff of 25 RPKM was used as previously described^15^. Maternal genes in 1-or 2-cell embryos were excluded from the analysis of 4- to 16-cell embryos. The dataset for 102-cell embryo was obtained from previous research^14^. A threshold of 25 TPM was set for gene expression based on the unique expression pattern of genes in E cells. Maternal loading genes were filtered using this cutoff, as evident by typical maternal genes oma-1, mex-5 et al. Eala and MSxppp cells were represented by 102-EX and 102-MSx in the plot. Genes in LADs were identified by at least 50% of the gene’s sequence lies within a 10 kb bin, or the gene spans the entire 10 kb bin. Gene position is based on ce11/WBcel235 genome annotations.

### Statistical analysis

All statistical analyses were performed using GraphPad Prism 10.

## Notes

### Competing Interest Statement

The authors have declared no competing interest.

