## Supplementary material for "Nuclear lamina-associated domain biogenesis is regulated by nuclear pore density during embryogenesis and mediates UV protection": Supp Figures

Supplementary Figure 1:

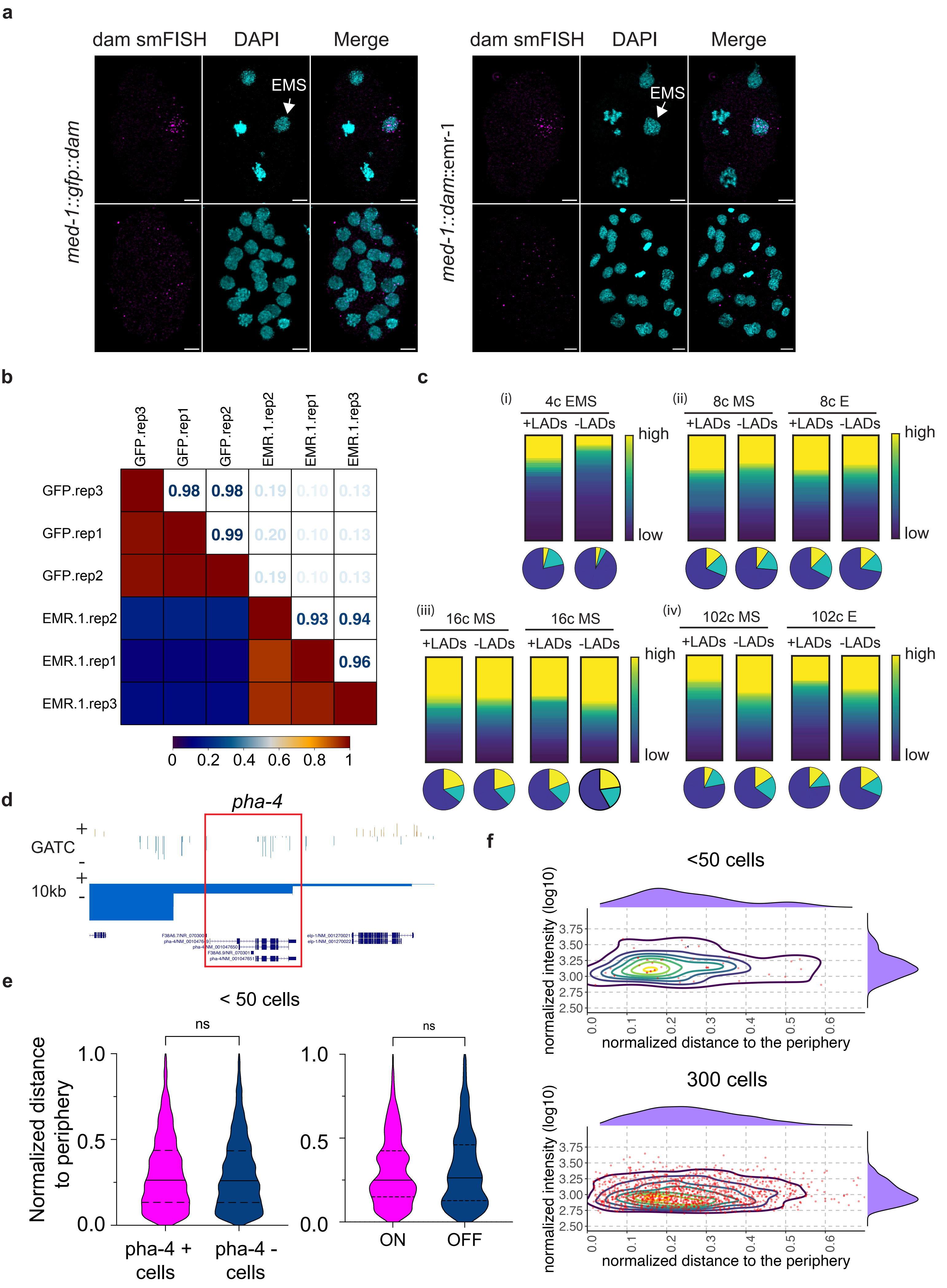

Supplementary Figure 2:

**a**

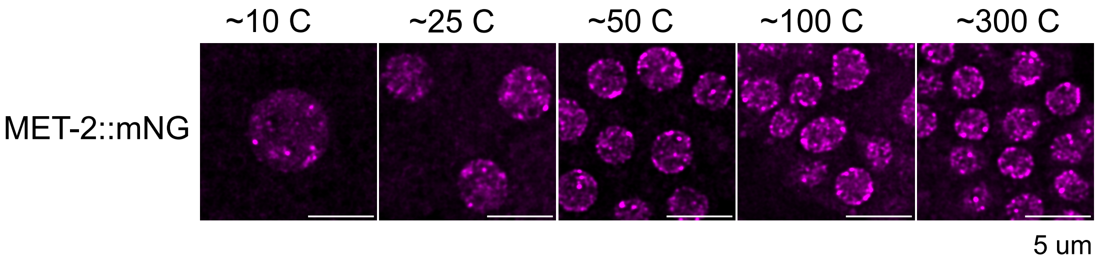


**b**

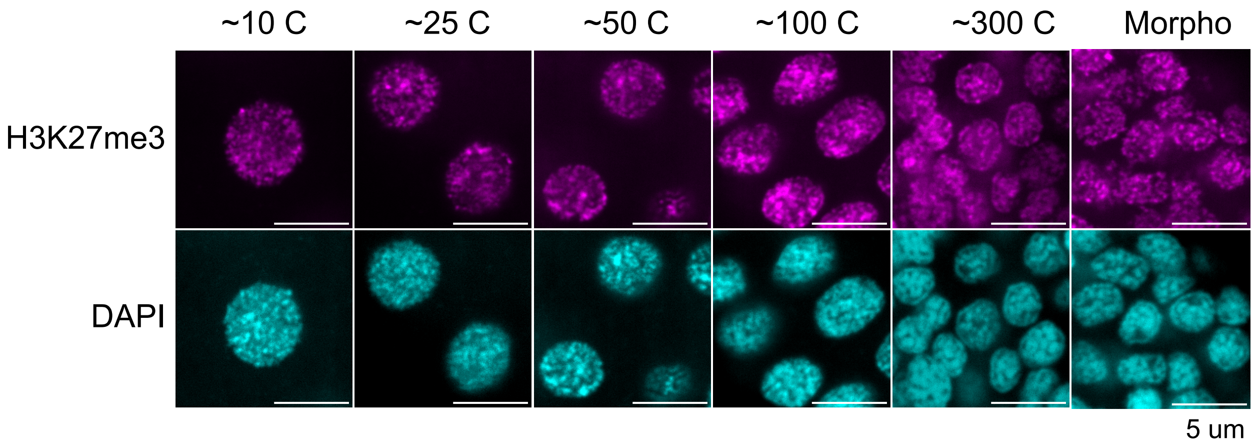


**c**

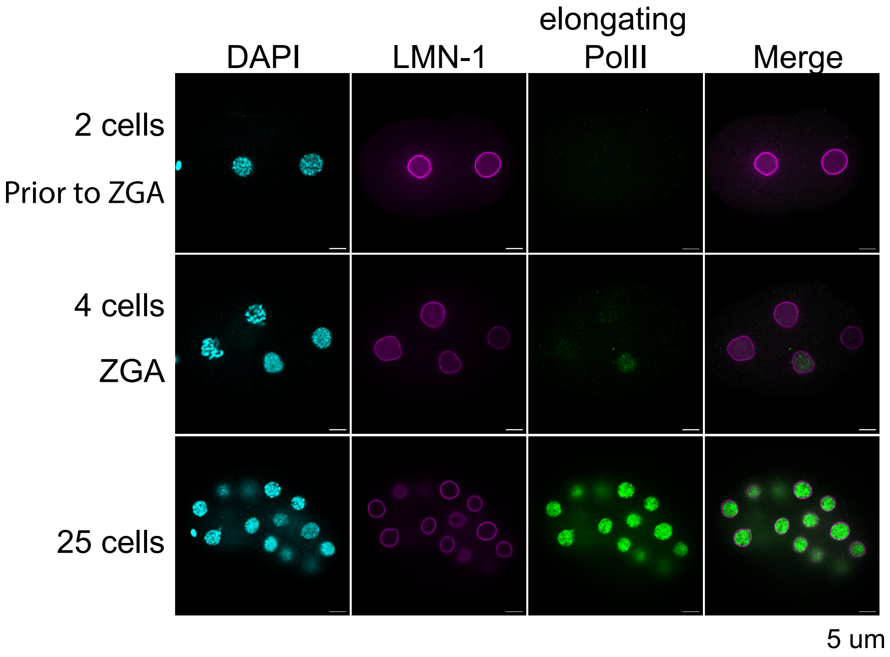

Supplementary Figure 3:

**a**

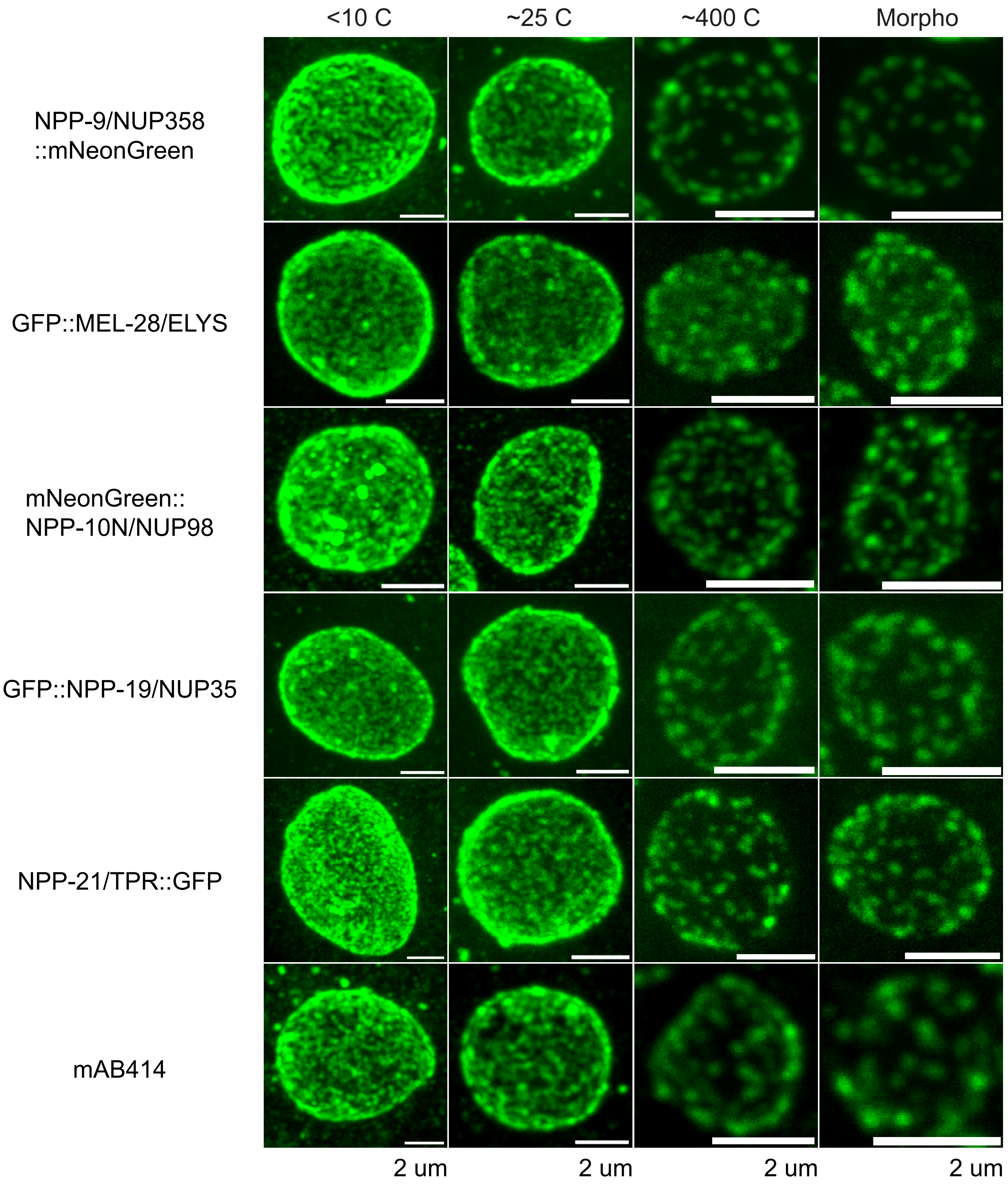

Supplementary Figure 4:

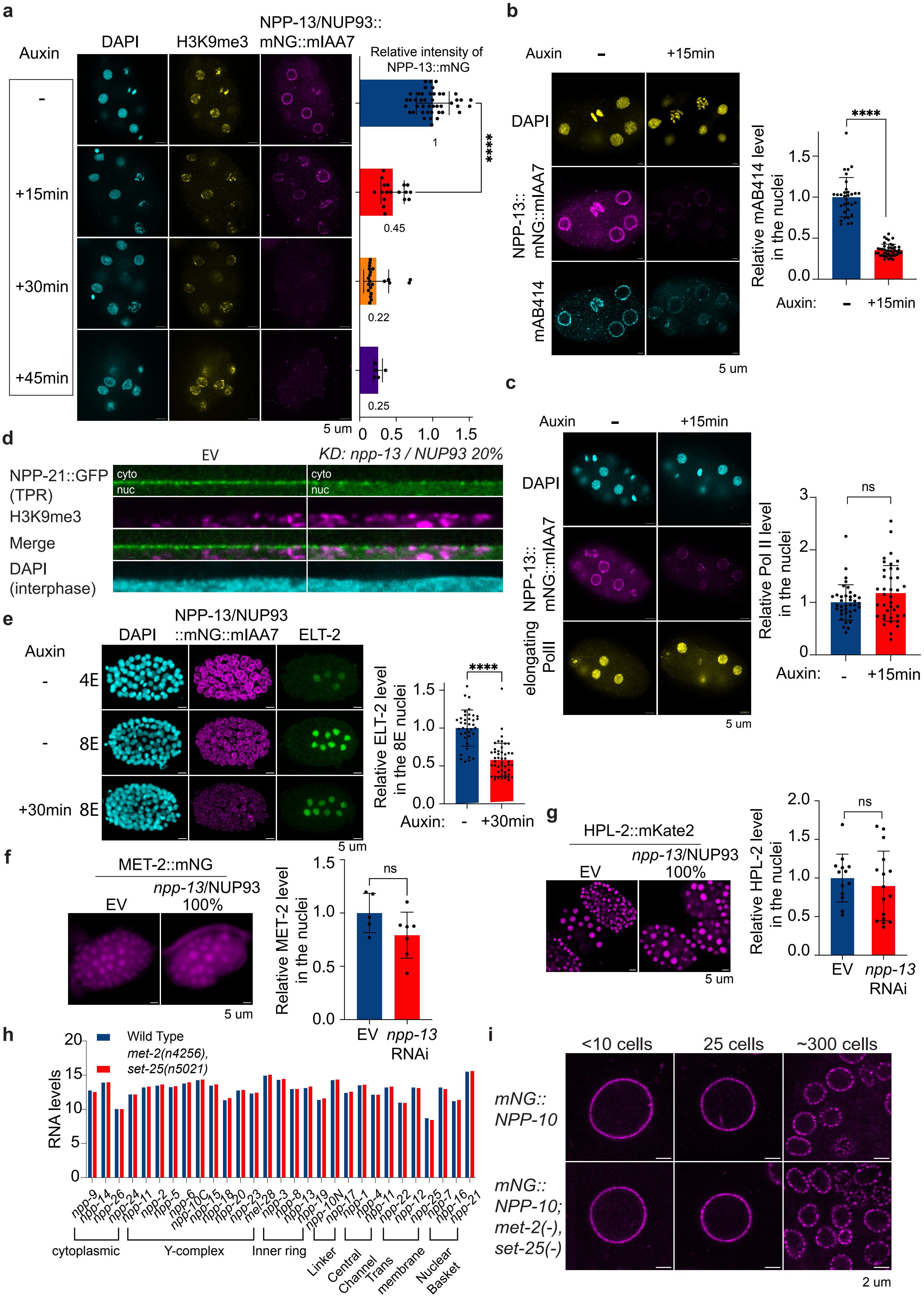

Supplementary Figure 5:

**a**

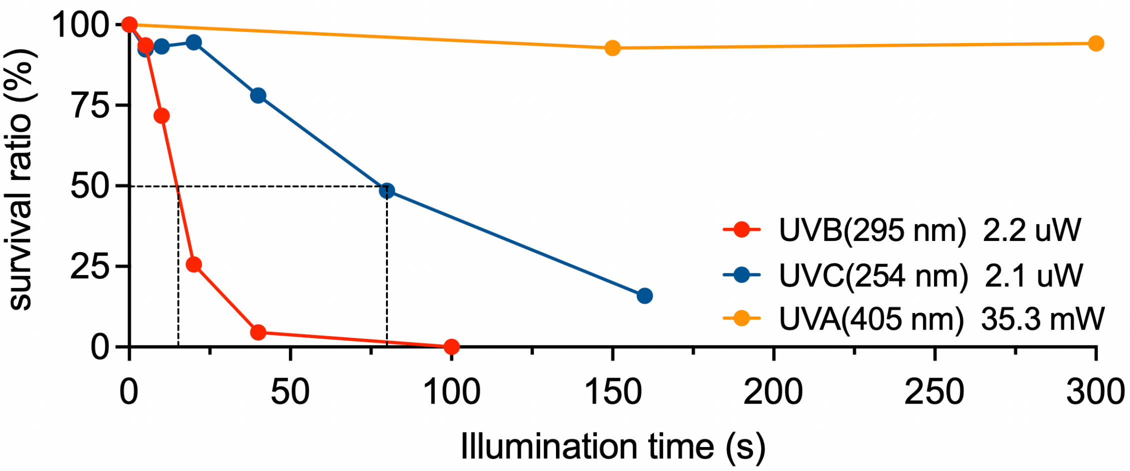
